# Encoding model for continuous motion-sensitive neurons in the intermediate and deep layers of pigeon Optic Tectum

**DOI:** 10.1101/2021.04.24.441242

**Authors:** Songwei Wang, Mengyu Zhao, Longlong Qian, Zhizhong Wang, Li Shi

## Abstract

There are typical neurons in the intermediate and deep layers of the optic tectum of avian, which are sensitive to small continuous moving targets. Based on this sensitivity of these neurons to continuous moving targets, the hypothesis of directed energy accumulation in dendrite field of these neurons is proposed. Based on the phenomenon that single dendrite activation can induce somatic spikes in vitro, the hypothesis of sequential probability activation mechanism of soma is proposed. Combined with the above hypotheses, the information encoding model of these typical neurons is constructed. Moreover, electrophysiological experiments and model simulations are carried out to obtain the response of neurons to visual stimuli of sequential motion, random motion and random sequential motion. Results show that the encoding model fits the response properties of continuous motion-sensitive neurons well. This study is of great significance for understanding the neurophysiological process of small-target perception for avian tectofugal pathway and the construction of the brain-inspired small-target detection algorithm.

## 1 Introduction

In avian, the majority of the cells in the intermediate and deep layers (800-1500μm) of optic tectum (OT) respond stronger to small moving objects than to stationary objects or wide-field motion (Frost & Difranco, 1976; Jassik-Gerschenfeld et al., 1975; Frost, 1993). Some of these motion-sensitive neurons receive direct synaptic input from retinal ganglion cells (RGC) at their distal dendritic endings (bottle-brush endings, bbe) in the superficial layers (Harald et al.,1998). When directly activating bbe (avoiding the inhibitory effect of synapses), it was found that as long as the time interval of continuous bbe activation was greater than a certain threshold (tens of milliseconds), a single activated bbe would induce somatic response with a high probability regardless of whether the positions of these activated bbe were the same (Luksch et al.,2004). Similar to these motion-sensitive neurons in avian, one synapse downstream of the retina, wide-field (WF) cells in the superficial superior colliculus (sSC) of mice (the physiologically corresponding nucleus of avian optic tectum in mammals) with the broad receptive fields respond comparatively strongly to small, slowly moving stimuli presented anywhere within a large region of space (Drager & Hubel,1975; O’Leary & McLaughlin,2005; Gale & Murphy,2014), but weakly (if at all) to large, suddenly appearing stimuli or full-field drifting gratings. Individual dendrites of WF cells can independently initiate strongly propagating dendritic spikes, and that nearly all spike output in response to visual stimuli is driven by such spikes (Endo et al.,2008; Gale & Murphy,2016).

That is, for the motion-sensitive neurons in avian and mice, a single dendrite is sufficient to drive the somatic response under certain conditions. In the classical neuron modeling, the somatic response is usually driven by the current obtained from the weighted sum of all dendritic inputs. Compared to the soma firing mechanism of classical neurons, the firing mechanism of motion-sensitive neurons appears more suitable for the detection of local small stimuli.

The preference for sequential continuous small moving targets is also an important feature of these motion-sensitive neurons. These neurons which react strongly to a continuously moving bar as well as an apparent motion stimulus, provided that the stimulus moved to a new position quickly and thus resembled the continuously moving bar (Verhaal & Luksch,2016).As for the WF cells of mice sSC, Gale’s results show that WF cells strongly preferred the long, continuous stimulus as long as the distance between successive locations was small, and the stimulus with local motion when the range of movement was larger (Gale & Murphy,2014).

Studies have shown that this preference of motion-sensitive neurons for small targets may stem in part from the synaptic inhibition of large horizontal cells that respond preferentially to the sudden appearance or rapid movement of large stimuli (Endo et al.,2003; Gale & Murphy,2016). Optogenetic reduction of their activity reduces movement selectivity and broadens size tuning in WF cells by increasing the relative strength of response to stimuli that appear suddenly or cover a large region of space (Gale & Murphy,2016). The horizontal cells in avian might constitute local inhibitory circuits within the retino-tectal synapses and, in addition, contribute to sensitivity to small and moving stimuli (Luksch & Golz,2003; Khanbabaie.et al.,2007).

Through the above literature, it can be found that there are a kind of continuous motion-sensitive neurons in the intermediate and deep layers of avian OT and mice SC which usually have sensitivity to small targets with continuous movement, and insensitivity to stationary objects or full-field stimulation. Their single dendrite has the ability to induce somatic response and the local inhibition from horizontal cells might contribute to their size selectivity. However, there is still no research on the encoding model of the neural mechanism of these neurons at present.

In this paper, based on the above advances, these following work is done to model the information processing of the continuous motion-sensitive neurons in the intermediate and deep layers of pigeon OT: (I) an information processing module of RGC-to-dendrite of continuous motion-sensitive neurons is presented to simulate the spike trains on the dendrite of this kind of neurons, and an inhibition mechanism is introduced to mediate the small target selectivity; (II) the key hypothesis of directed energy accumulation is given to explain the preference to continuous motion; (III) a sequential probabilistic activation hypothesis (SPA) is proposed to simulate the ability of single dendrite to induce somatic response. Moreover, the corresponding electrophysiological experiments were carried out, and the effective fitting of the model response to the neurons’ response was obtained under different stimuli, which verified the effectiveness of the proposed model. This study could furthermore deepen the understanding of the neurophysiological process of small-target perception for avian and is of great significance for the construction of the brain-inspired target detection algorithm.

## 2 Materials and methods

### Animals, surgery and electrode implantation

#### Animals

Twenty pigeons (*Columba livia*) weighing 300-400g were obtained from a local dealer (Gongchuang Pigeon, Henan, China) and housed in an animal facility for at least 2 weeks prior to the experiments. The animals were maintained under a 12-h light: dark cycle at a constant temperature of 23 ± 2°C with free access to water and food.

#### Surgery

Each pigeon was anesthetized with sodium pentobarbital (3%, 0.17 mL 100 g^−1^) and placed in a stereotaxic apparatus (model ST-5ND-B, Chengdu Instrument, Chengdu, China). A small window was opened to expose the left lateral tectum. The right eye was held open and the left eye was covered. Multi-unit activity was recorded with a multi-electrode array comprising 16 polyimide-insulated platinum/iridium microwires (Clunbury Scientific, Bloomfield Hills, MI, USA), which were arranged in 2 rows with 8 wires in each row (electrode diameter = 50μm; electrode spacing = 350μm; row spacing = 350μm; impedance = 20-50kΩ). The array was lowered approximately 800-1500μm below the tectum surface using a micromanipulator (Mc1000e, Siskiyou, San Diego, CA, USA). A silver wire from the array was connected to a bone screw inserted into the skull surrounding the surgical openings for grounding. The experimenter intermittently monitored the anesthetized pigeons to assess their eye movements during data recording and no eye movements were observed.

#### Ethics statement

All of the experimental procedures conformed to the institutional guidelines for the care and use of laboratory animals at Zhengzhou University, Henan, China and the National Institutes of Health Guide for Care and Use of Laboratory Animals (GB 14925-2001). The Ethics Review Committee of Zhengzhou University of Life Sciences approved this paper. Pigeons were killed at the end of the experiment with sodium pentobarbital (3%, 0.50mL 100 g^−1^).

### Stimulus protocols

Visual stimuli were generated using a PC running a Matlab toolbox (Psychtoolbox; MathWorks, Natick, MA, USA) and displayed on a CRT monitor positioned 40 cm in front of each pigeon’s right eye. The screen had a refresh rate of 100 Hz, a spatial resolution of 1080×1080 pixels. The screen grayscale values ranged from 0 to 255. Three grayscale values(0, 128, 255 corresponding to black, gray and white respectively) were mainly used in these experiments and would not be changed unless stated.Before recording, the horizontal axis of each pigeon’s head was rotated by 38° so the lateral fovea of the right eye was contralateral to the exposed tectum and given a projected screen size of 32.4°×32.4°.

### Identification of cell types and Determination of receptive field

Here, movement point stimulus, sparse noise stimulus and ON/OFF stimulus were designed and displayed sequentially to find whether these collected neurons are sensitive to continuous movement and insensitive to sudden appearance of random targets and change of brightness (wide-field stimulus). These three kinds of stimuli were displayed on the full screen and the corresponding ranges of receptive fields were determined by the intensity response of these neurons (DeAngelis et al.,1995).

#### Movement point stimulus

First, movement point stimulus was used (Fig. 1a) to find these neurons which have relatively large receptive field of motion based on the sensitivity to small continuous moving objects. On the gray background, a black target point, 0.9°×0.9°, was moved continuously at 0°, 90°, 180° or 270° with a velocity of 90°/s, starting and ending at different positions on the edge of the background until all rows or columns on the background had been passed once for every direction. The same forms of motion were repeated 5 times for each trial. The receptive field of motion was determined as the field with the highest activity, whose boundaries were the positions where the activity fell below 25% of the maximum response.

**Fig. 1.**
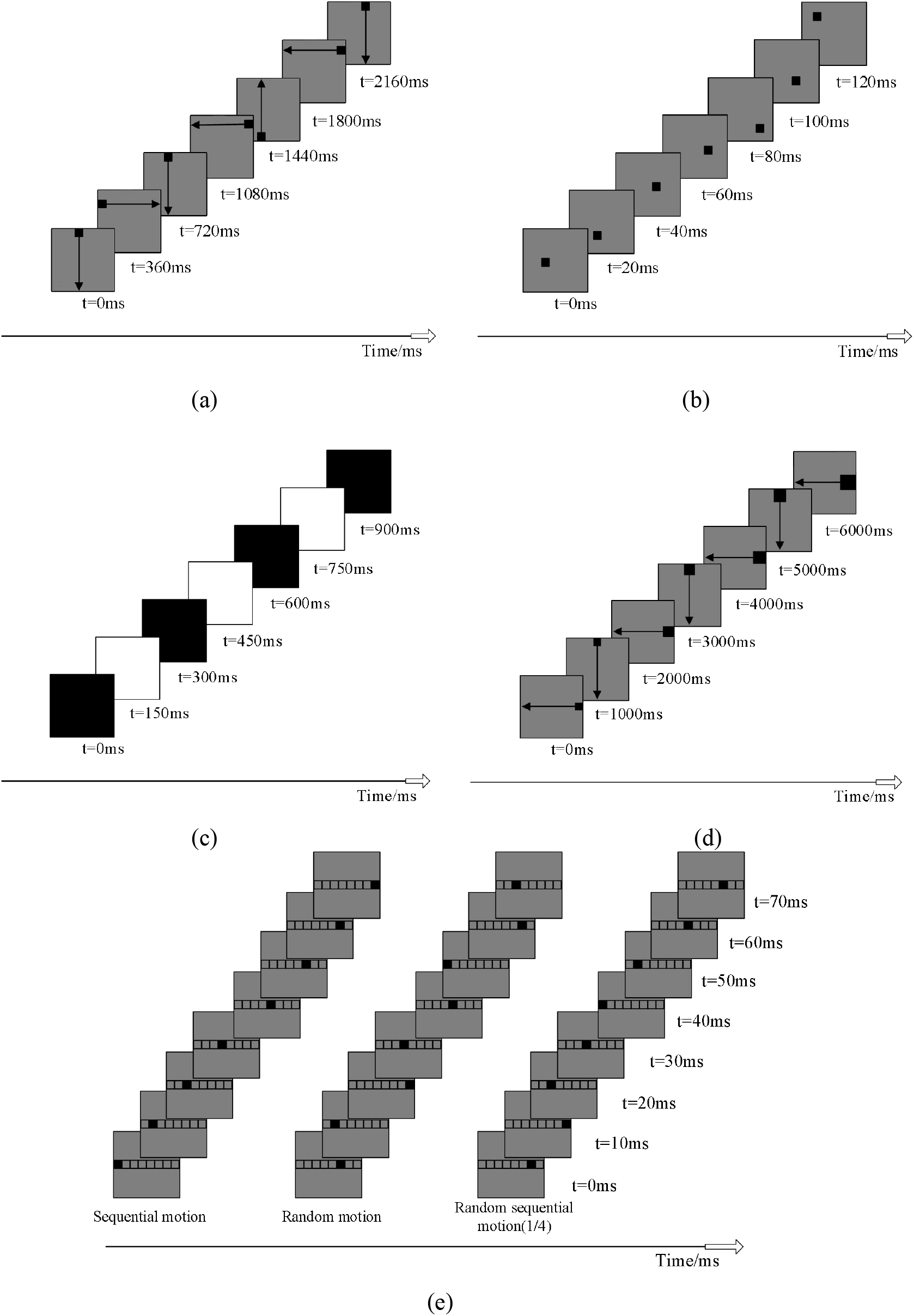
Stimulus protocols. (a) Movement point stimulus. (b) Sparse noise stimulus. (c) ON/OFF stimulus. Multi-size stimulus. (e) Continuous sensitivity stimuli. Random sequential motion (1/8) didn’t shown in here. The black target points in these figures are not shown with the real target size, just a hint. Note that movement point stimulus, sparse noise stimulus and ON/OFF stimulus were displayed on the full screen, but multi-size stimulus and continuous sensitivity stimuli were displayed only on the receptive field. The real sizes of background images in the example of (a), (b) and (c) are 32.4°×32.4°. The size of receptive field in the example of (d) were 12°×12°. The black arrows in (a) and (d) indicate the direction of the target movement.

#### Sparse noise stimulus

The randomly blinking gray-black checkerboard stimulus mode, which was also called sparse noise stimulus, was carried out to further determine whether the neurons found by the previous stimulus were sensitive to the sudden appearance of random targets. A single black grid (0.9°×0.9°) over a gray checkerboard was presented in different locations of the screen (36×36 grid) for 20ms. The sequences of these locations were repeated 20 times following different pseudorandom orders, as shown in Fig. 1b.

#### ON/OFF stimulus

The simple black and white luminance stimulus model (i.e. ON/OFF stimulus) was used to study the response characteristics of this kind of neurons to a light switching on and off (wide-field stimulus) as shown in Fig. 1c. Black and white stimulation were repeated alternately and each of them was presented for 150ms. The number of repetitions was 80 for each trial.

### Verification of key properties

Multi-size stimulus and continuous sensitivity stimuli were designed to verify the tolerant extent for these continuous motion-sensitive neurons found by above stimuli to the small target and continuous motion. Note that these two kinds of stimuli were just displayed on the corresponding receptive field of neurons.

#### Multi-size stimulus

In the multi-size stimulus which was used to test the size sensitivity, the black target points of different sizes (0.36°×0.36°, 0.72°×0.72°, 1.8°×1.8°, 3°×3°, 7.5°×7.5°) were traveled through the center of the receptive field with a speed of 12°/s, starting and ending at the edge of the receptive field. This stimulus repeated 10 times for each size as shown in Fig. 1d.

#### Continuous sensitivity stimuli

Sequential motion, in which a black target point progressed from one adjacent location to the next, random motion, in which the point appeared at the same locations but in random order and random sequential motion, similar to sequential movement, in which the point moved locally near each random position, were designed and executed to verify the preference for continuity and our hypotheses as shown in Fig. 1e. There were two types of random sequential motion: 1/4 and 1/8 (didn’t shown in Fig. 1e) which mean the rate of the path length of the local sequential motion to the total path length. The speed of the black target point was 12°/s. The size of the black target point is the preferred size of the corresponding neuron. Each group was repeated 10 times for each trial.

### Action potential detection and pretreatment

The signals were collected using a Cerebus system (Blackrock Microsystems, Salt Lake City, UT, USA) and amplified 4000 times. The spikes were first extracted by band-pass filtering (second-order Butterworth) the raw signals between 250 Hz and 5 kHz at a sampling rate of 30 kHz, and detected with a threshold set at a signal-to-noise ratio of 1.50. The spiking events were saved for offline analyses, which were performed using MATLAB.

### Encoding model for continuous motion-sensitive neurons

The following parts present our encoding model in detail. Firstly, the dendrite field of this kind of continuous motion-sensitive neurons is modeled, whose dendritic endings are found to be cosine distribution about the center of the dendritic field (Luksch et al.,2006). Considering they receive inputs from individual RGC unit and synaptic inhibition from horizontal cells, the simulated information processing module from RGC to dendritic endings was constructed which mainly refer Songwei Wang’s work (Songwei Wang et al.,2019) and an effective spatial center-surround antagonism was introduced into this module to mediate the small target selectivity. Then, the soma of continuous motion-sensitive neuron may receive multiple spike trains from activated dendrites induced by targets. Since continuous motion-sensitive neurons prefer long continuous motion, the key hypothesis of directed energy accumulation is given to explain this characteristic and smoothly the sequential probability activation hypothesis of soma is proposed to explain the ability to induce somatic response by their single dendrite.

### Dendrite field modeling of continuous motion-sensitive neurons

Luksch has provided a method for modeling the dendritic field of a class of motion-sensitive neurons which have somata in stratum griseum centrale (SGC), extend their sparse dendritic fields with cosine distribution and make monosynaptic contact with axon terminals from retinal ganglion cells (RGCs) (Karten et al.,1997; Luksch et al.,1998; Mahani et al.,2006). This method was adopted to model the dendritic field of continuous motion-sensitive neurons in this paper, whose dendritic endings were cosine distributed about the center of the dendritic field:

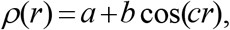

where fitted parameters *a, b* and c could be tuned manually based on the size of receptive field.

The inhomogeneous “Poisson cluster” process was used to generate the simulated dendritic fields of this kind of neurons, which has been shown to work well in (Diggle, 2003; Mahani et al.,2006; Khanbabaie et al.,2007). In the Possion cluster process, the probability of having a parent location is equal to *ρ*(*r*)/2×ΔA for any small area Δ*A* of the dendritic field, which means it depends on the distance, *r*, of that small area from the center of the dendritic field of a neuron. Centered on each parent location, then, two dendritic endings were generated out of a symmetric 2D Gaussian distribution. The probability of having a parent location in a small area Δ*A* is independent of other small areas and is not uniform across space for a continuous motion-sensitive neuron.

### Integration mechanism of continuous motion-sensitive neurons dendrites on input visual information

The dendrites of each motion-sensitive neuron receive the output of spike trains of RGC neurons (Frost,1993; Karten et al.,1997; Luksch et al.,1998; Mahani et al.,2006; Luksch & Golz,2003). Since the difficulty of collecting the signal from RGC axon, this paper adopts the method used by Songwei Wang (Songwei Wang et al.,2019) to simulate the signal that transfer to the dendrite of continuous motion-sensitive neuron. In (Songwei Wang et al.,2019), by analyzing the STC (spike-trigger covariance), the ON filter and OFF filter of RGC could be deduced based on the response of the superficial and intermediate OT neurons which are sensitive to the change of light to particular designed stimuli. This method was directly used in our paper to get the ON/OFF filters of RGC (Fig. 2a). Then, the simulated signal transfer from the RGC axon to the dendrite of continuous motion-sensitive neuron could be expressed as Fig. 2b which includes linear filtering, half-wave rectification, sum and thresholding.

**Fig. 2.**
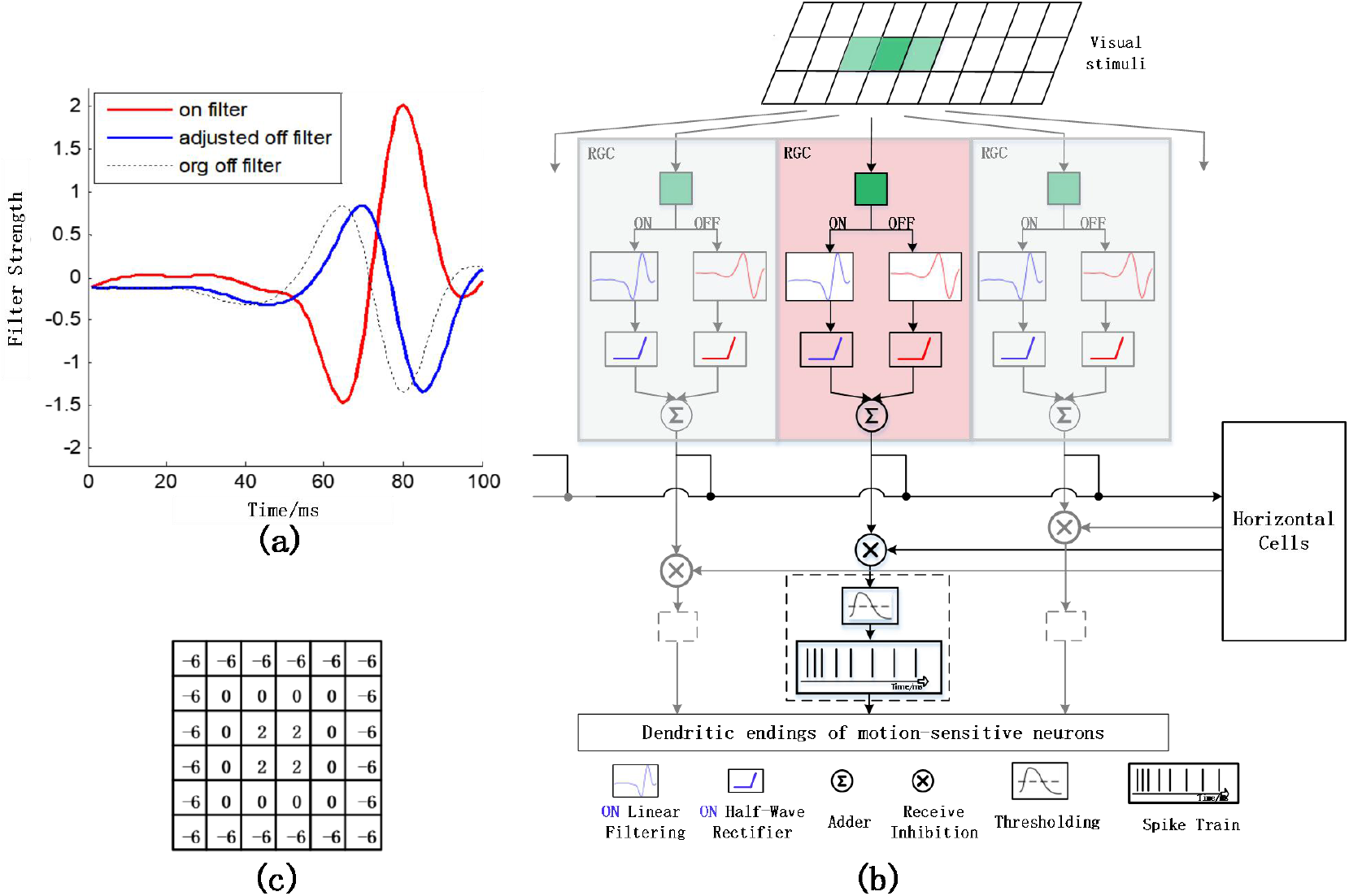
The establishment of integration mechanism on input visual information. (a) The ON/OFF filters of RGC that used in this paper. (b) The simulated information processing module from RGC to dendritic endings of continuous motion-sensitive neurons. Visual stimuli are filtered, rectified, added-up by RGC and inhibited by horizontal cells, and finally converted into dendritic spike trains of continuous motion-sensitive neurons. (c) The kernel that are used in spatial centre-surround antagonism.

Whereas continuous motion-sensitive neurons responded most strongly to relatively small and continuously moving stimuli, horizontal cells, which have long and horizontally extending dendrites with relatively sparse branching, responded best to the sudden appearance or brisk motion of larger stimuli and they receive monosynaptic retinal inputs on their dendritic endings which are usually in the same layer as the motion-sensitive neurons’ (Endo T et al.,2003; Luksch & Golz,2003; Khanbabaie et al.,2007; Gale & Murphy,2014; Gale & Murphy,2016). One possible explanation for the opposing selectivity of visual response of continuous motion-sensitive neurons and horizontal cells is that horizontal cells integrate the inputs of multiple RGC and produce inhibition effects on the dendritic endings of continuous motion-sensitive neurons as shown in Fig. 2b. To simulate the inhibition effects, the strong spatial centre-surround antagonism as shown in Fig. 2c was added to the process of simulated signal transfer. Target size tuning is achieved by varying the gain and spatial extent of this centre-surround antagonism.

### Modeling the continuous motion-sensitive neurons ‘ preference for continuous motion

For a continuous motion-sensitive neuron, ideally, a continuous moving target passing through its receptive field will activate a series of dendrites, and the connecting line of these activated dendrites can roughly reflect the target track. Continuous motion-sensitive neurons prefer small targets with long duration of continuous motion (Gale & Murphy,2014; Verhaal & Luksch,2016). To model this special response attribute, the following assumptions were given:

#### Directed energy accumulation hypothesis

After a certain dendrite **s1** is activated at time *t*, it will generate a facilitative area R1 in its vicinity that could last a certain period of time (Fig. 3a) and obtain an energy E1. This area is a predicted area where the dendrite **s1** predicts future activated dendrites by targets would appear in.

**Fig.3.**
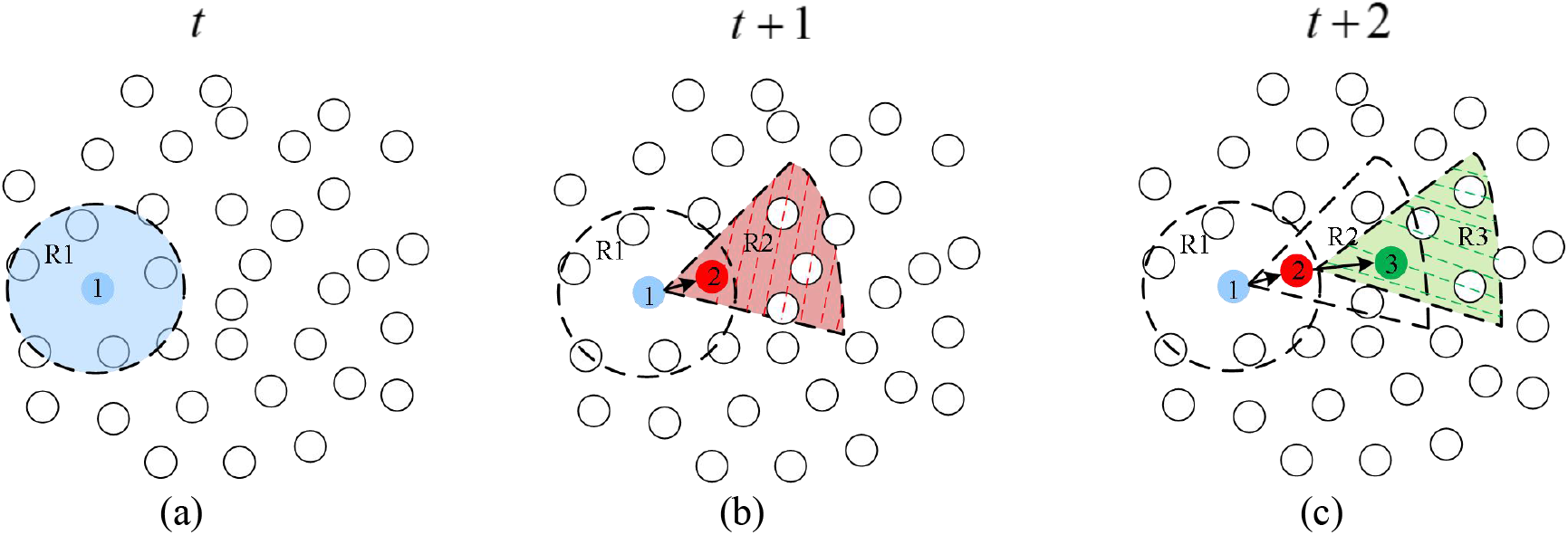
Diagram of directed energy accumulation hypothesis. The establishment of integration mechanism on input visual hypothesis. The small black circles represent dendrites of a continuous motion-sensitive neuron. (a) At time *t*, dendrite 1 (blue little circle) is activated and forms a facilitative area R1 (the blue circle area) that could last a certain period of time. (b) At time *t* + 1, dendrite 2 (red little circle) is activated and appears in R1, forming a track Γ (1->2) with dendrite 1 and a facilitative area R2 (the red sector). (c) At time *t* + 2, dendrite 3 is activated and appears in R2, forming a track Γ (1->2->3), and a facilitative area R3.

If an activated dendrite **s2** appears in the facilitative area R1 of dendrite **s1** before the disappearance of this area at time *t* +1, dendrite **s1** and dendrite **s2** could form a track Γ and dendrite **s2** will generate a “stronger” directed facilitative area R2 along the track Γ that also could last a certain period (Fig. 3b). To reflect this directed continuous activation effect, dendrite **s2** will obtain an energy E2 that equal to the sum of energy E1 and energy 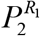 which denotes the probability of the current activated dendrite **s2** falling into the previously facilitative area R1. In other words, dendrite **s2** accumulates energy E1 and energy 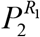 contributed by itself.

At time *t* + 2, if a dendrite **s3** is activated within the facilitative area R2 of dendrite **s2** before the disappearance of this area, correspondingly, there will be a longer track Γ, a higher energy E3, and a facilitative area R3 as shown in Fig. 3c. If the dendrite is activated without the facilitative area *R*_2_ (it’s not shown in Fig. 3), it will initiatively accumulate energy like dendrite **s1**.

#### Formal description of the directed energy accumulation hypothesis

Suppose that for a continuous motion-sensitive neuron who have *N* dendrites, the positions of these dendrites are fixed. At time *t*, there is a dendrite activated by a small moving target which can be represented as a 2-dimensional vector

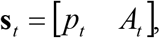

where *p*_*t*_ represents the position of this dendrite, and *A*_*t*_ represents the activate intensity of this dendrite, namely, the generated spike train.

Then, **s**_*t* +1_ and **s**_*t* +2_ at time *t* +1 and *t* + 2 are activated respectively in the same way as shown in Fig. 4. Since most moving targets in nature move continuously, it can be considered that the velocity (including magnitude and direction) of the target remains almost constant when it passes through the dendrite field of the continuous motion-sensitive neuron. Correspondingly, the small moving target will continuously activate several dendrites. Due to the irregularity of the dendrite field and the maneuverability (generally weak, but also existing) of the real physical motion target at short time, the central connecting line τ(the track formed by black arrows as shown in Fig. 4) of these continuously activated dendrites **s**_*t*_, **s**_*t* +1_ and **s**_*t* +2_ is usually not a straight line, but a broken line.

**Fig.4.**
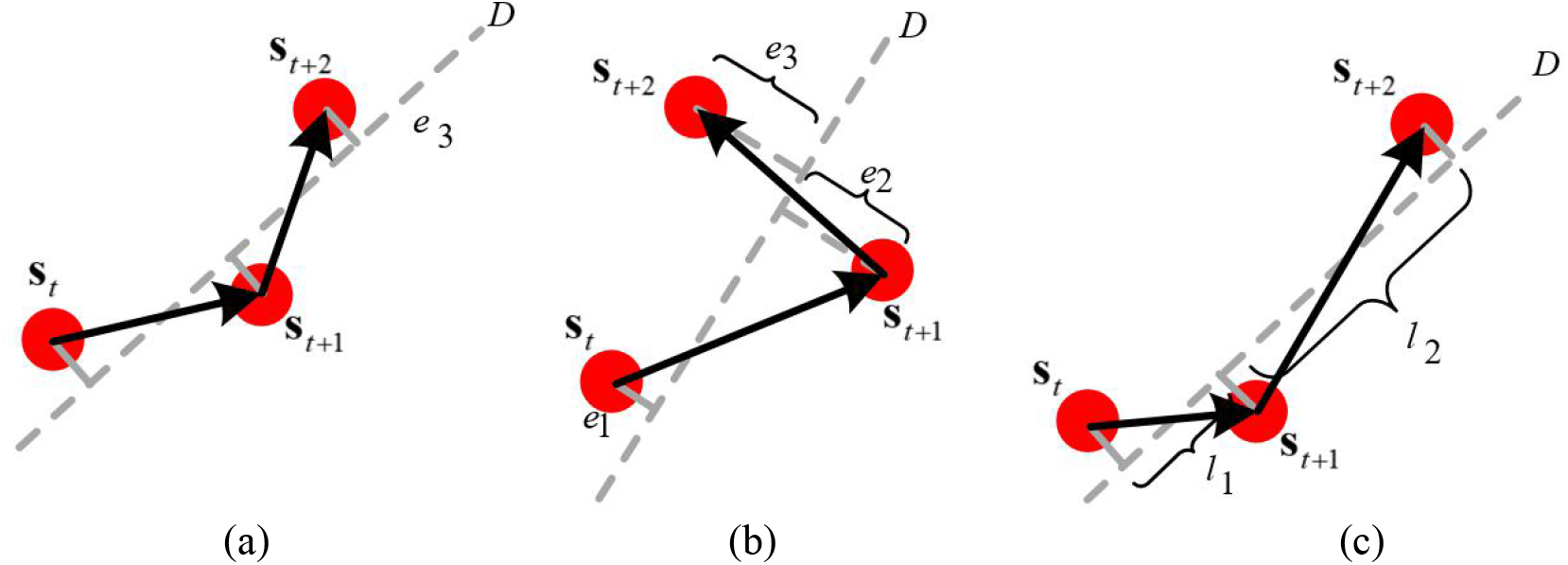
Judgment of target track. (a) The three sequentially activated dendrites meet the range of linearity and velocity fluctuation, that is, the broken line τ formed by black arrows is one possible target track Γ. (b) The three dendrites do not meet the range of linearity fluctuation, so **s**_*t* +2_ does not belong to the facilitative region of **s**_*t* +1_. (c) The three dendrites do not meet the range of velocity fluctuation, that is, **s**_*t* +2_ does not belong to the facilitative region of **s**_*t* +1_.

Considering the weak maneuverability of the targets and the preference for continuous motion of continuous motion-sensitive neurons, if the linearity and velocity fluctuation degree of the broken line τ less than the “broken” threshold, it is considered the broken line τ is a target track Γ as shown in Fig. 4a, which means all activated dendrites along Γ have fallen into the facilitative areas of the previously activated dendrites. On the contrary, if the linearity or velocity fluctuation degree of the broken line τ is greater than the “broken” threshold, the **s**_*t* +2_ is unable to accumulate the previous energy E and correspondingly the broken line τ is excluded from the target tracks in Fig. 4b and 4c.

According to the position information *p*_*t*_, *p*_*t* +1_ and *p*_*t* +2_ of the three dendrites, the straight line *D* fitted by them could be obtained, and the distances *e*_*t*_, *e*_*t* +1_ and *e*_*t*+2_ between the line *D* and the three dendrites respectively can be estimated as shown in Fig. 4b. The linearity *S*_*τ*_ of τ can be formulate as:

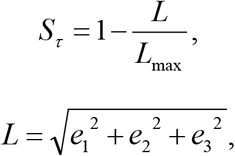

where *L* is the linearity fluctuation and *L*_max_ is the maximum linearity fluctuation threshold.

Furthermore, by calculating the projected distance *l*_1_ between sequential activated dendrite position *p*_*t*_ and *p*_*t* +1_ on line *D*, and the projected distance *l*_2_ between sequential activated dendrite position *p*_*t* +1_ and *p*_*t* +2_ on line *D* as shown in Fig. 4c. The velocity fluctuation degree *S*_*vec*_ of τ can be formulate as:

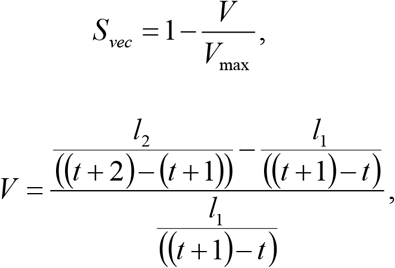

where *V* is the velocity fluctuation and *V* _max_ is the maximum velocity fluctuation threshold.

The probability *P* that the current activated dendrite falls into the facilitative area of the previously activated dendrite can be defined as

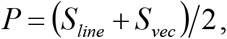

For dendrite **s**_*t* +2_, the energy it accumulated can be calculated as follows:

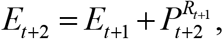

Where 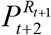 is the probability that the activated dendrite **s**_*t* +2_ falls into the nearby

facilitative region *R*_*t* +1_ of a certain dendrite **s**_*t* +1_ whose accumulated energy is *E*_*t* +1_.

As initialization, when the first dendrite of the continuous motion-sensitive neuron is activated at time *t*, its accumulated energy is

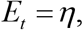

where *η* is a constant.

### Modeling the ability of single dendrite to induce somatic response

The classical information integration mechanism of soma of neuron is weighted sum, mathematically, which is equivalent to the inner product operation, namely the measurement of similarity. The more similar the input image pattern is to the internal template of neurons, the stronger the response of neurons will be. Therefore, classical neurons can be considered as feature extractors as shown in Fig. 5a. However, the single dendrite of continuous motion-sensitive neuron is found to have the ability to activate the soma, which means that they are suitable for the detection of local small stimuli.

**Fig.5.**
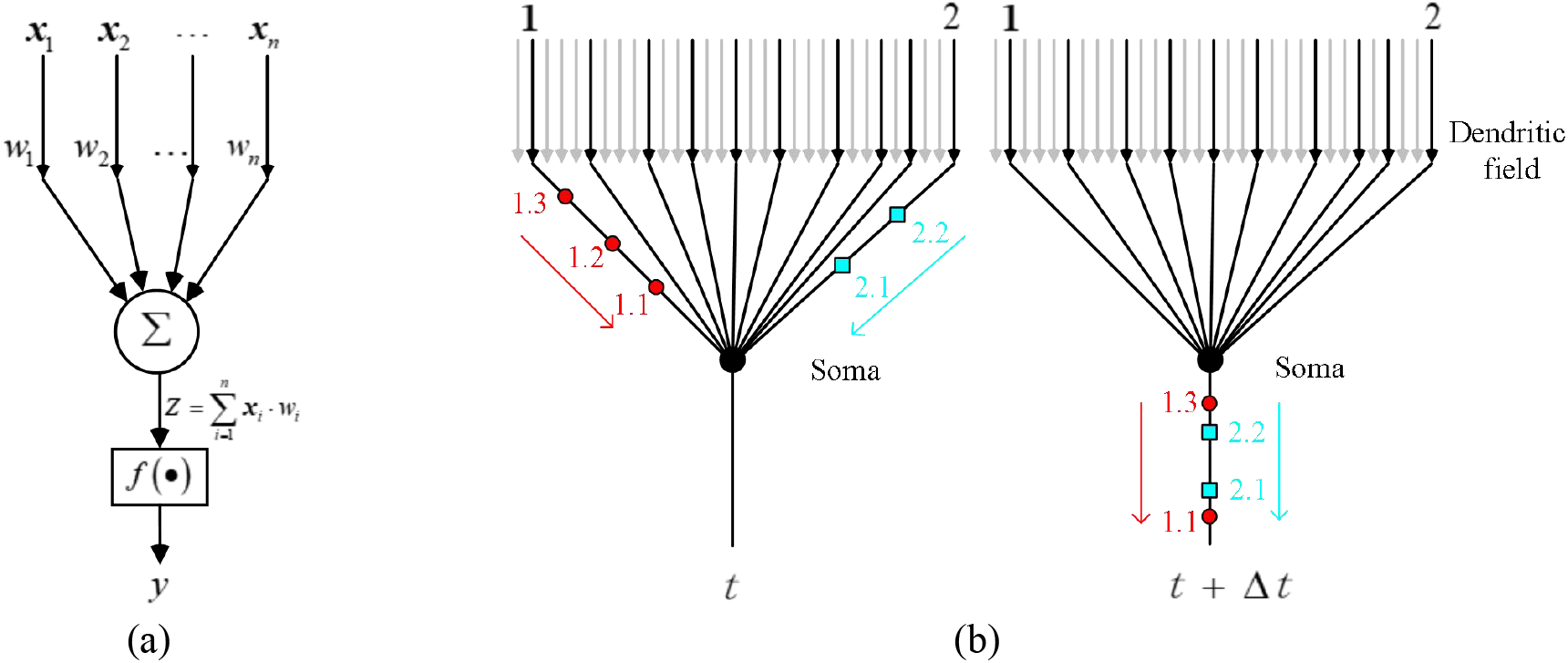
The different information integration mechanism of soma. (a) The classical information integration mechanism of soma of neurons is weighted sum. Each component of the input vector is multiplied by its corresponding weight and summed, and then the nonlinear transformation is performed. (b) Dendrite 1 and dendrite 2 are activated at time *t* and generate a series of spike trains respectively. 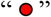 represents the spike trains generated by dendrite 1, 1.1,1.2 and 1.3 represent the serial number of each spikes respectively. 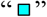 represents the spike trains generated by dendrite 2, 2.1 and 2.2 represent the serial numbers of each spike respectively. According to the SPA hypothesis, the spike trains generated by these two dendrites will flow through the soma in sequence with a certain probability that determined by the accumulated energy themself. As shown in figure of *t* + Δ*t* moment, the spikelet sequence 1.1 and 1.3 of dendrite 1 flowed through the soma, but 1.2 did not pass, while the spikelet sequence 2.1 and 2.2 of dendrite 2 both flowed through the soma.

Here, the hypothesis of sequential probability activation (SPA) mechanism of soma of continuous motion-sensitive neurons is proposed to explain this characteristic: the ability of dendrite to induce somatic response is determined by the accumulated energy itself. That’s mean if one dendrite **s**_*t*_ of continuous motion-sensitive neuron is activated and produces a series of spike trains at time *t*, these spike trains will flow through the soma (induce somatic response) in sequence with a certain probability which depends on the level of accumulated energy *E*_*t*_. The greater the energy, the more possibility these spike trains can flow through. As for multiple dendrites, each dendritic spike train activates the soma according to the respective accumulated energy as shown in Fig. 5b.

## 3 Result

### The model dendritic field of continuous motion-sensitive neurons

It has long been hypothesized that structure and function are correlated in neural systems (Stuart et al.,1999; Jan & Jan,2003). The simulated dendritic field of continuous motion-sensitive neurons can be generated according to the above method whose dendritic endings are cosine distributed about the center of the dendritic field. Moreover, the distribution of nearest neighbor distances was fitted by an exponential function between pairs of nearest neighbor dendritic endings from the one continuous motion-sensitive neuron irrespective of their absolute distance from the center of the dendritic field (Luksch et al.,2006). Here a simulated dendritic field was symbolically enumerate where fitted parameters *a* = 1.01# /100 × 450*μm*^2^, *b* = 1.04# /100× 450*μm*^2^ and c = 0.0016 rad/*μ*m^2^ as shown in Fig. 6a and the corresponding fitted exponential curve was obtained using the simulated dendrite fields with the same parameters as shown in Fig. 6b.

**Fig.6.**
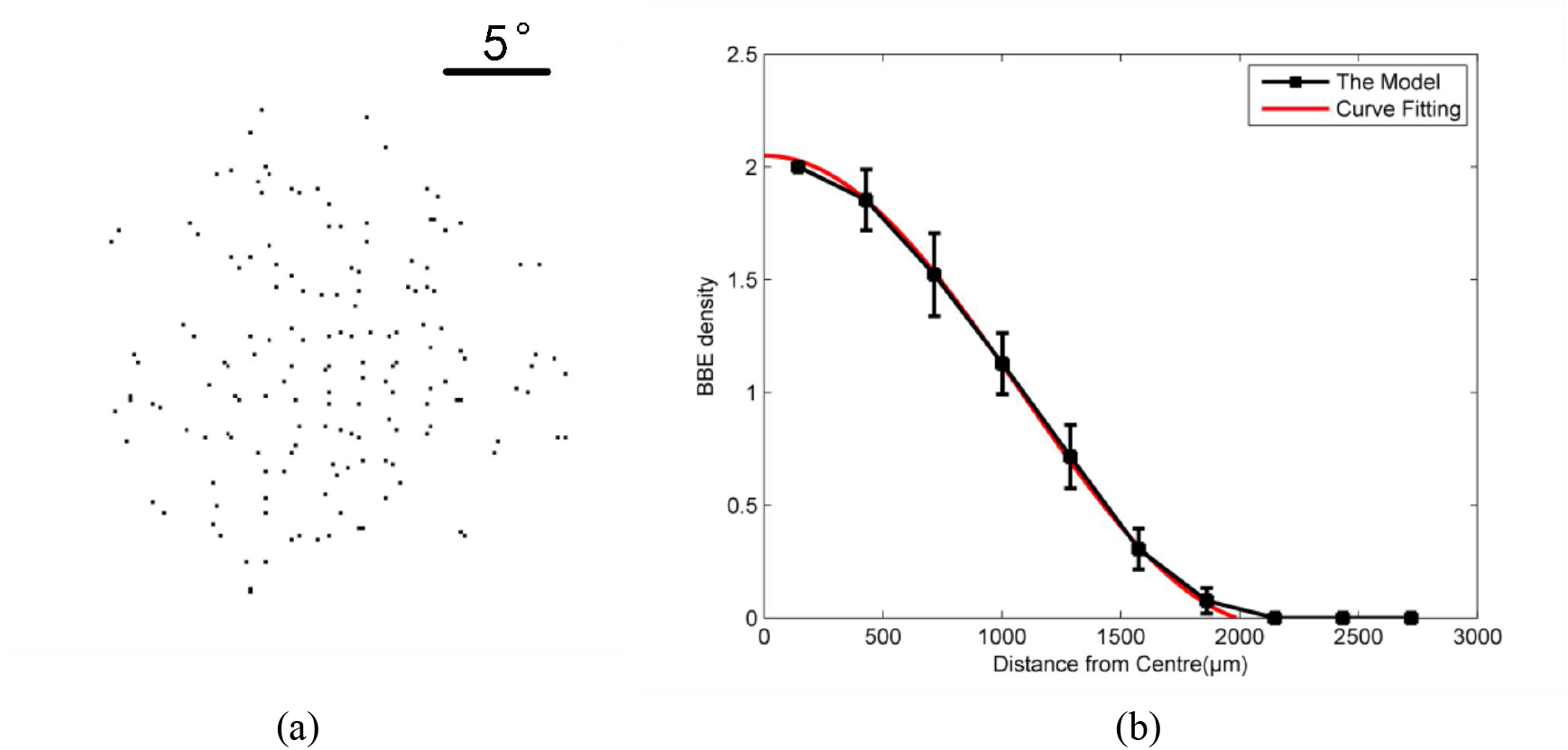
The model result of dendritic field. (a) Simulated inhomogeneous Poisson cluster spatial distribution dendritic endings in a circular dendritic field with cosine density function. (b) Average density, ρ(r), of “bottle brush” dendritic endings (bbe) per continuous motion-sensitive neuron as a function of the distance from the center of the dendritic field. The density was measured from 100 continuous motion-sensitive neurons. Error bars are standard error. The red line was obtained by fitting a cosine function of the form ρ(r) = a +b cos (cr).

### Comparison of receptive field of motion and sparse noise receptive field

The data in this paper are collected from the response signals of continuous motion-sensitive neurons in the intermediate and deep OT area (800-1500μm central gray layer) of 16-channel-microelectrode carrier pigeons. First of all, the spatial central positions and receptive fields of these collected neurons were obtained by reverse correlation (Deangelis et al.,1995) according to the response of neurons to movement point stimulus and sparse noise stimulus. Receptive fields of motion and sparse checkerboard receptive fields help us rule out those neurons that are not continuous motion-sensitive neurons. Several continuous motion-sensitive neurons that have different sizes of receptive fields are shown in Fig. 7 (Neuron 1,Neuron 2 and Neuron 3). Furthermore, based on the insensitivity of continuous motion-sensitive neurons to large targets or wide-field motion, these neurons who had no sensitivity to on-stimulus and off-stimulus were identified as continuous motion-sensitive neurons (Fig. 7. Neuron 1, Neuron 2 and Neuron 3). Here, a total of 14 neurons from 20 pigeons were identified as continuous motion-sensitive neurons. In addition, another kind of motion-sensitive neurons that are sensitive to continuous movement, sudden appearance of random targets and change of brightness (wide-field stimulus) were found as shown in Fig. 7 (*Neuron 4*).

**Fig.7.**
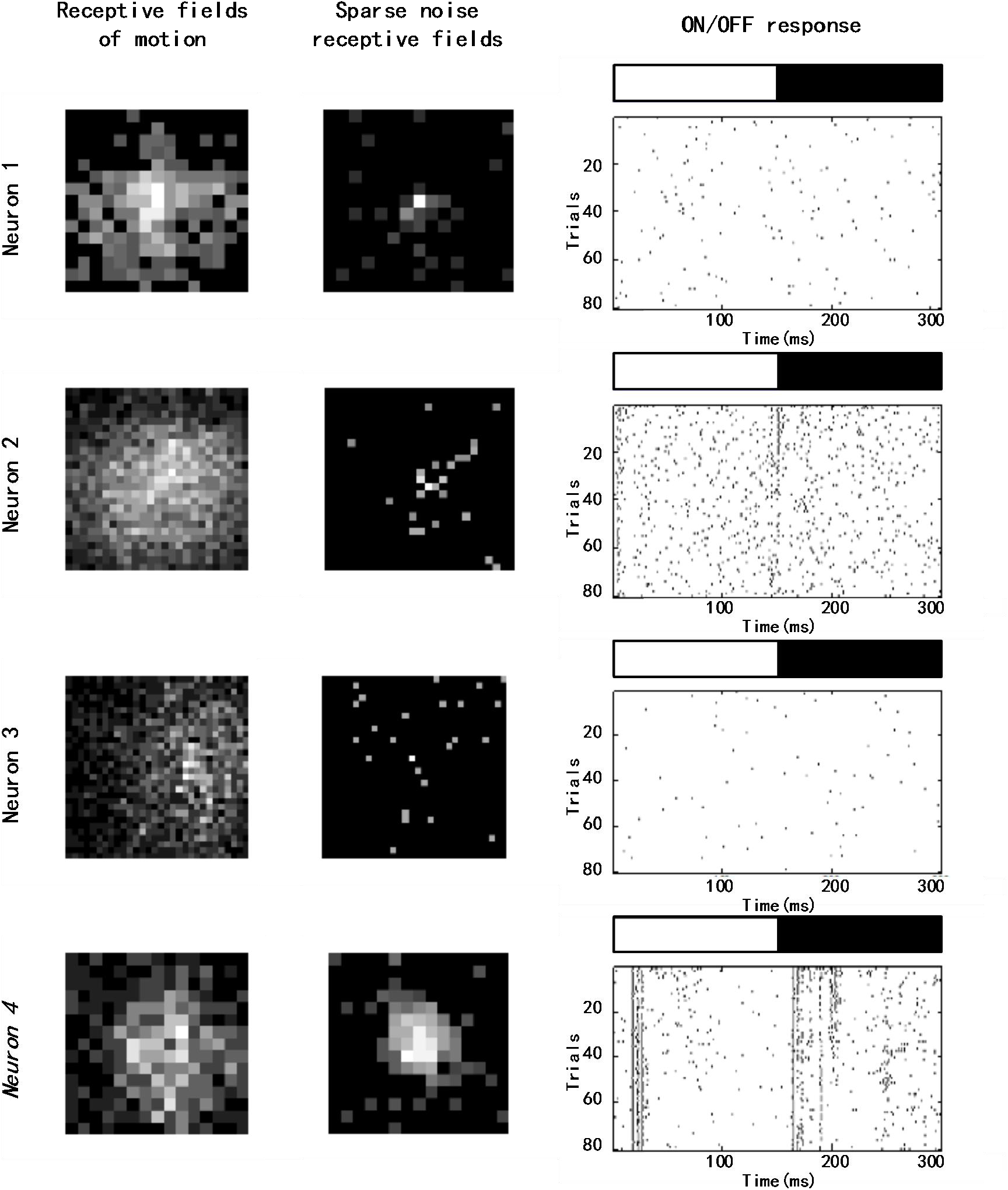
Response of motion-sensitive neurons to movement point stimulus, sparse noise stimulus and ON/OFF stimulus. The size of every grid in receptive field of motion and sparse receptive field is 30×30 pixels. The sizes of receptive fields (Neuron 1, Neuron 2, Neuron 3, *Neuron 4*) were 13.5°×13.5°,22.5° ×22.5°, 27°×27° and 13.5°×13.5°. Neuron 1, Neuron 2 and Neuron 3 were continuous motion-sensitive neurons that we need. *Neuron 4* is one another kind of motion-sensitive neuron that we found accidentally.

### Target size tuning

After finding these continuous motion-sensitive neurons, the size tuning property was first tested using multi-size stimulus. Here the response results of two neurons in electrophysiological experiments were presented as shown in Fig. 8, and it was apparently observed that the response of continuous motion-sensitive neurons was the strongest when the target crossed the center of the dendrite field, which response trend is consistent with the response trends of Gale and Luksch (a spot moving through the center of the receptive field) (Gale & Murphy,2014; Verhaal & Luksch, 2016).The firing numbers under different sizes were counted and the size tuning property is presented as shown in Fig. 9 (red line). These continuous motion-sensitive neurons preferred the small moving targets about 0.72°×0.72° in size according to the size tuning. Considering that the response results of every neuron at the optimal size could be fixed to a uniform length of time through interpolation and sampling, the statistical response result of electrophysiological experiments under the respective optimal size could be gotten as shown in Fig. 10a.

**Fig.8.**
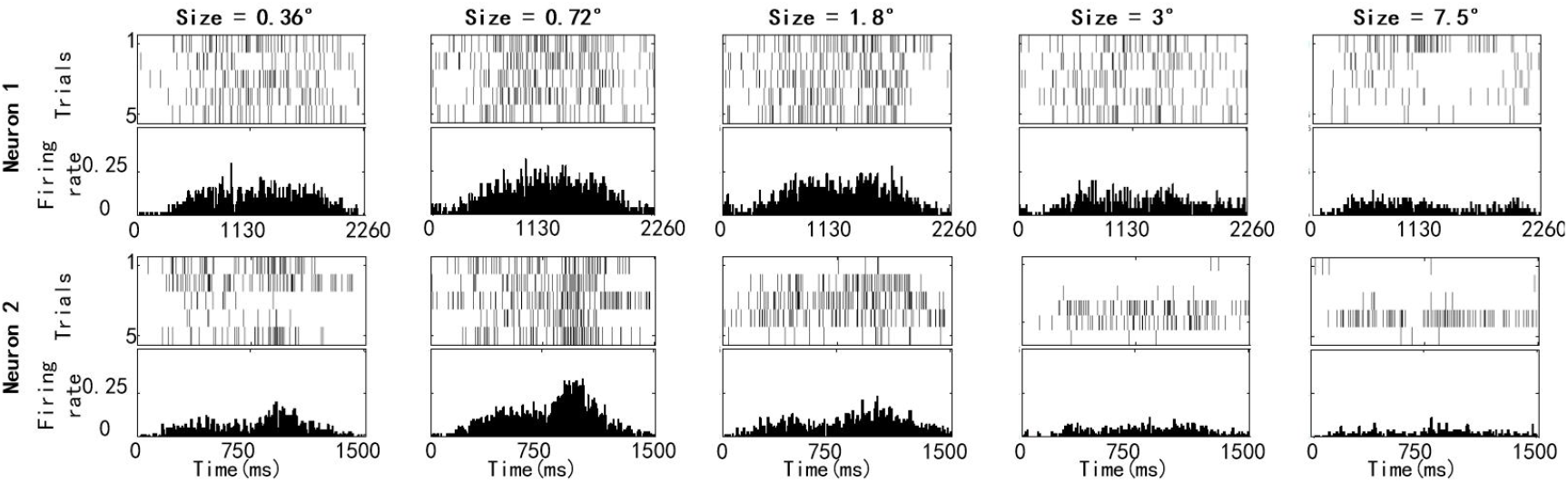
Raster diagram and mean firing rate of continuous motion-sensitive neurons with different sizes of continuous moving targets. Each motion under different sizes repeated 10 times. The sizes of receptive fields (Neuron 1, Neuron 2) are 27°×27° and 18°×18°.

**Fig.9.**
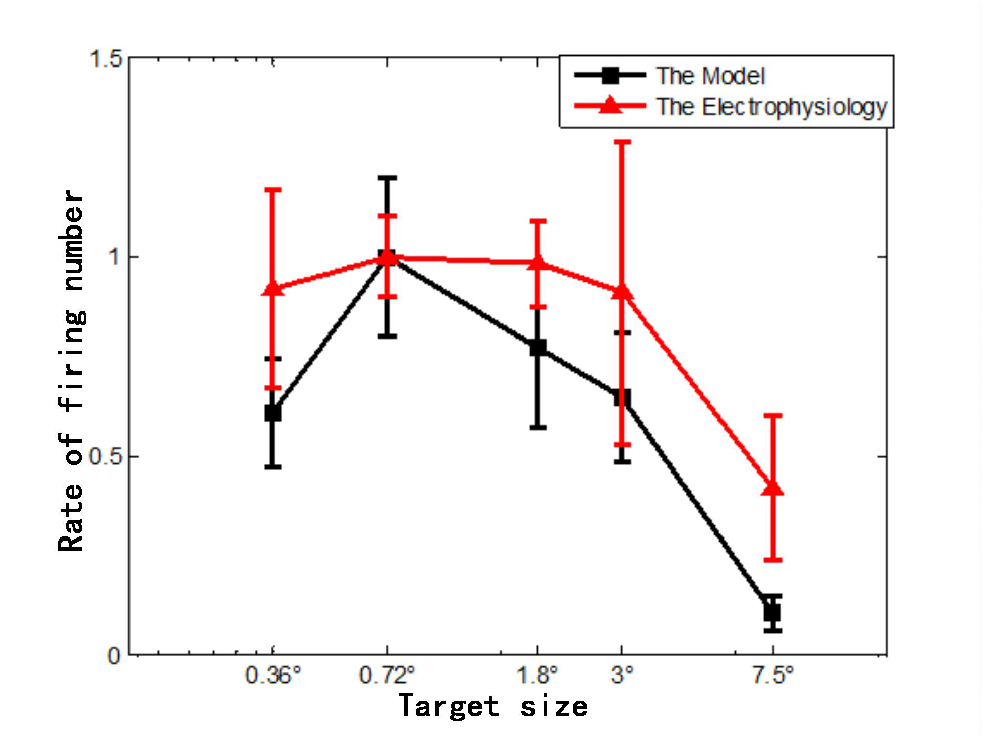
The tuning curves of continuous motion-sensitive neurons to multi-size stimulus. The red triangle line represents the result produced by the electrophysiology using 14 collected continuous motion-sensitive neurons. The black square line represents the result produced by model under using 300 simulated continuous motion-sensitive neurons. The firing number of each neuron for each size was calculated, and then divided by the firing number of its optimal preference size to normalize. The rate of firing number of all neurons for every size was then added up and normalized again. The horizontal axis is the logarithmic axis and represents the sizes of the small target, and the vertical axis represents the mean value and variance of rate of firing number.

**Fig.10.**
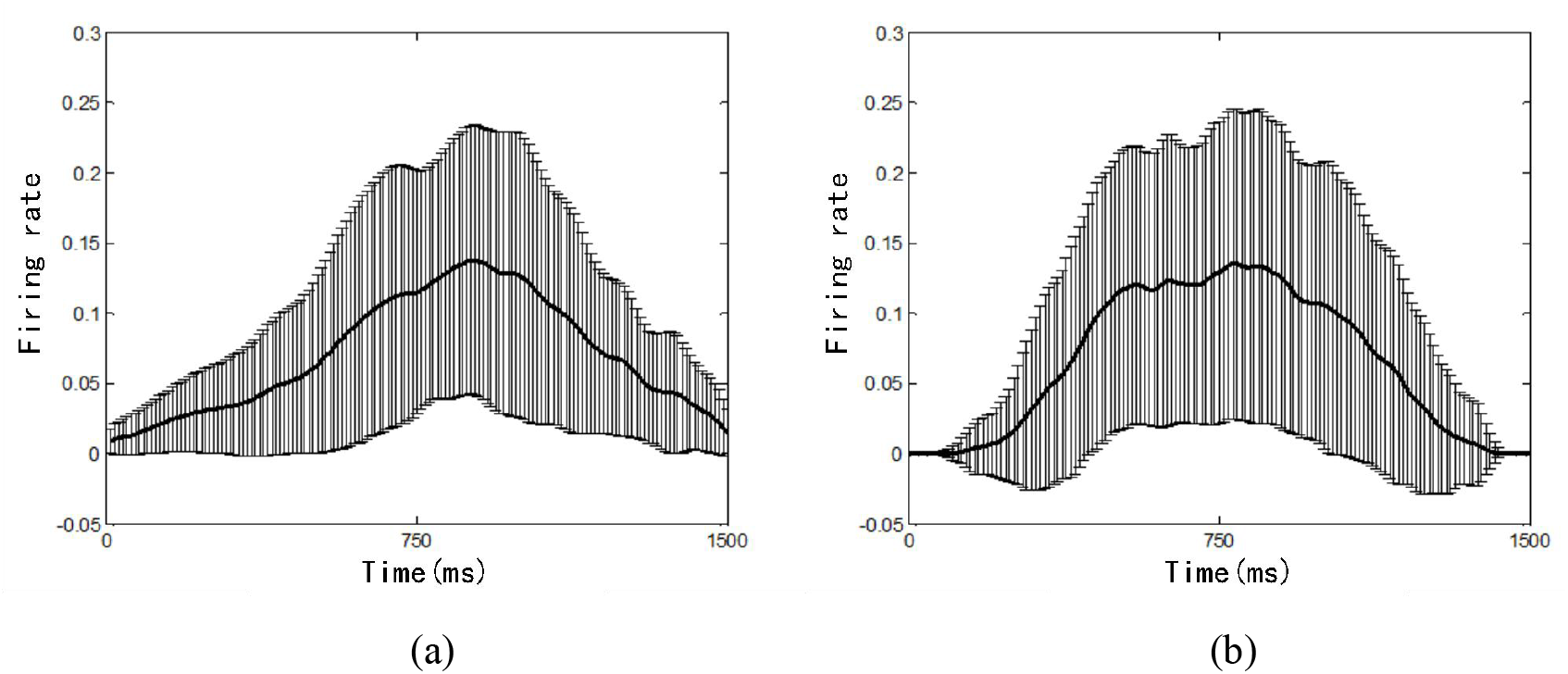
The statistical response result under the optimal size. The thick solid line represents the mean value of firing rate, and the thin solid line represents its corresponding variance. The average curves were smoothed by Gauss filter. (a) The electrophysiological experiment result using 14 collected continuous motion-sensitive neurons. (b) The model result. To be more statistical, 300 simulated continuous motion-sensitive neurons were randomly generated and counted which had somas located in one same center of the receptive field.

As for the response results of model under the same multi-size stimulus, it was found that these results are similar to the “bell curve” of electrophysiological experiment results. Here, the size tuning property and the statistical response result of model which had receptive field (18°×18°) was shown in Fig. 9 (black line) and Fig. 10b. Although the statistical firing rate curve of electrophysiological experiments was sharper than that of model, but their overall response trend was consistent. Using the hypotheses presented above, why the “bell curve” appeared could be explained. First, the RGC receives and processes the stimulus signal at different moments, then passes it (receiving the inhibitory effect of horizontal cells) on to dendritic endings of continuous motion-sensitive neuron. When the target starts to enter the receptive field at the initial period, some dendrites will be activated and carry a series of dendritic spike trains. Due to the relatively sparsity of dendrites at the edge of dendritic field, it is hard for these activated dendrites to form effective tracks Γ and accumulate enough energy *E* to completely induce somatic response, which resulting in weak response at the initial period. When the target moves near the center of the dendrite field, the dendrites in the center area of the dendrite field are relatively dense, which leads more effective tracks Γ and relatively large dendritic energy *E*, so that the soma response induced by the activated dendrites is strong. Then, the sparse dendrites at the edge area of the dendrite field gradually lose the accumulated energy as the moving target gradually leaves the dendrite field, that resulting in the weaken somatic response gradually. The similarity of the size tuning curves between electrophysiological experiments and model illustrate the validity of the integration mechanism of dendrites and hypotheses we proposed.

### Test for sensitivity to continuous motion

A remarkable characteristic of continuous motion-sensitive neurons is that they respond more to continuous moving targets than to stationary ones (Gale & Murphy,2014). Three kinds of stimuli, sequential motion, random motion and random sequential motion were tested in electrophysiological experiments and model to explore whether continuous motion-sensitive neurons are sensitive to the continuous or discrete moving targets. Random sequential movement indicates the degree of preference of continuous motion-sensitive neurons to target motion continuity. The longer the path length of the local sequential motion is, the more continuous the track is. The electrophysiological response of three continuous motion-sensitive neurons were shown in Fig.11 (Neuron 1, Neuron 2 and Neuron 3). It is obvious that they strongly preferred the sequential motion stimulus than to other stimuli. The response of the random sequential motion gradually became larger as the rate of the path length of the local sequential motion to the total path length increased and the bell curve response could be found in these response. As for the random motion, the response of continuous motion-sensitive neurons was consistently weak. In addition, the response of one another kind of motion-sensitive neuron, which was mentioned above, was also shown in Fig. 11 (*Neuron 4*). It was also sensitive to random motion beside sequential motion, but when the target was moving continuously, its response became a “bell curve” as the response of continuous motion-sensitive neurons.

**Fig.11.**
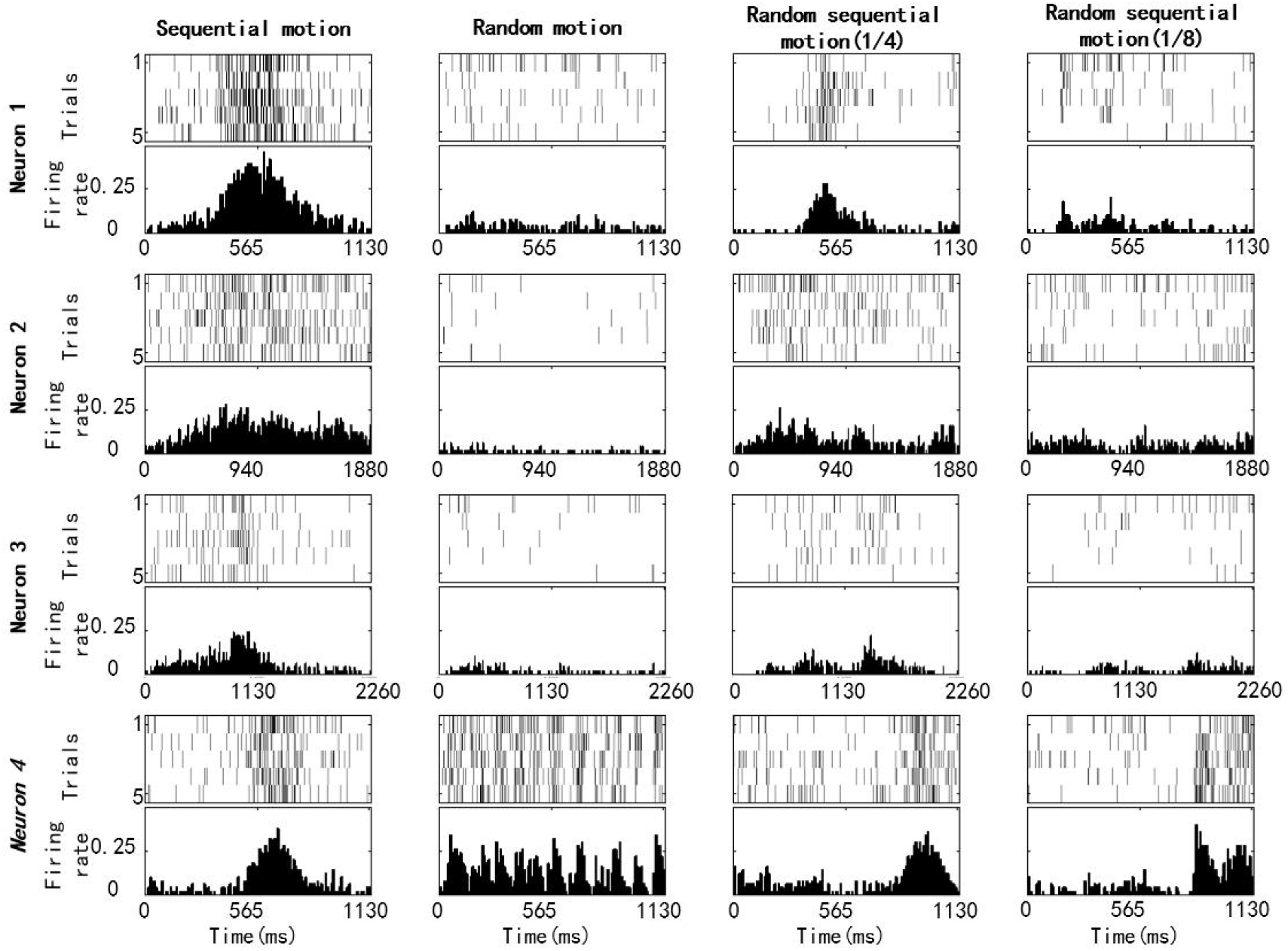
The electrophysiological results under three kinds of stimuli. Each motion repeated 10 times. The sizes of receptive fields (Neuron 1, Neuron 2, Neuron 3, *Neuron 4*) were 13.5°×13.5°, 22.5°×22.5°, 27°×27° and 13.5°×13.5°. Neuron 1, Neuron 2 and Neuron 3 were continuous motion-sensitive neurons that we need. *Neuron 4* was one anther kind of motion-sensitive neuron that we found accidentally.

The firing numbers under different continuous sensitivity stimuli were counted and the tuning curves for these three forms of motion is presented as shown in Fig.13 (red line). It is clear that continuous motion-sensitive neurons prefer motion with high continuity. In the same way, the statistical response result of electrophysiological experiments to continuous sensitivity stimuli could be gotten as shown in Fig. 12a and 12c (except that the response statistical response result to sequential motion because of the similarity to Fig. 10a) through interpolation and sampling.

**Fig.12.**
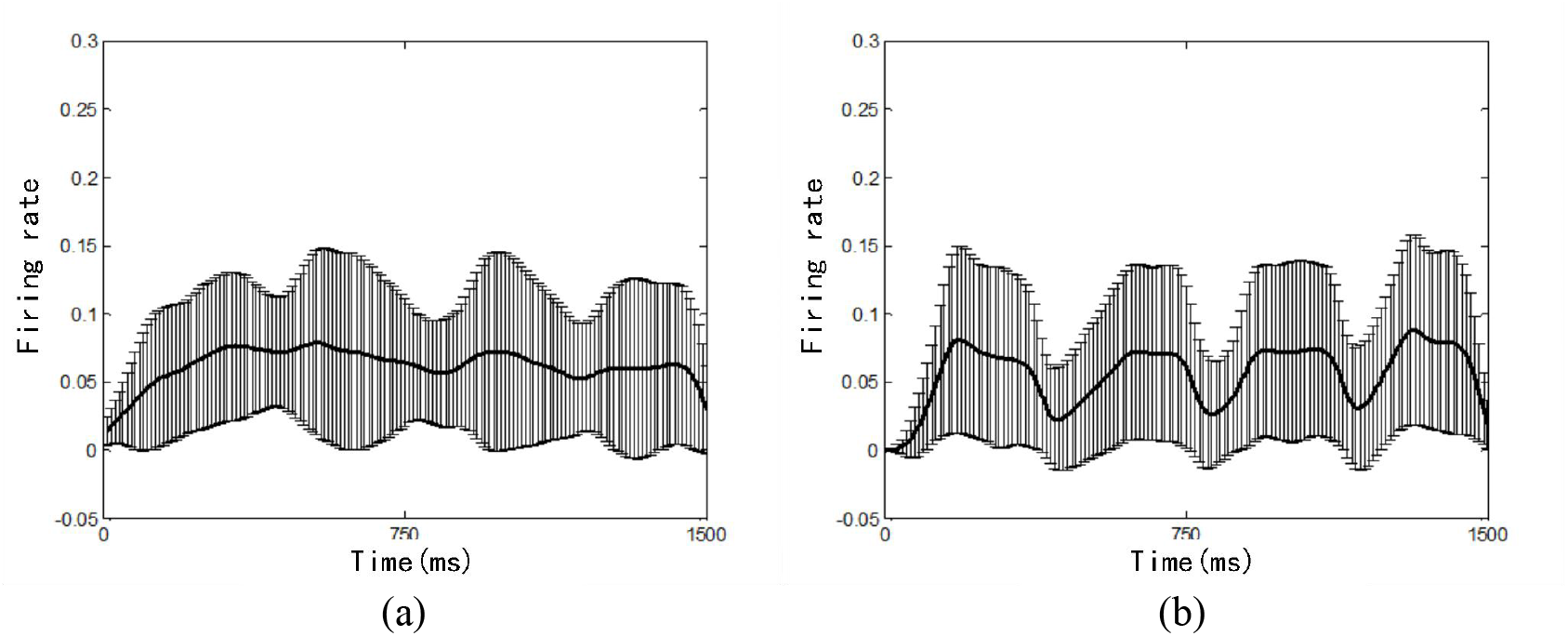
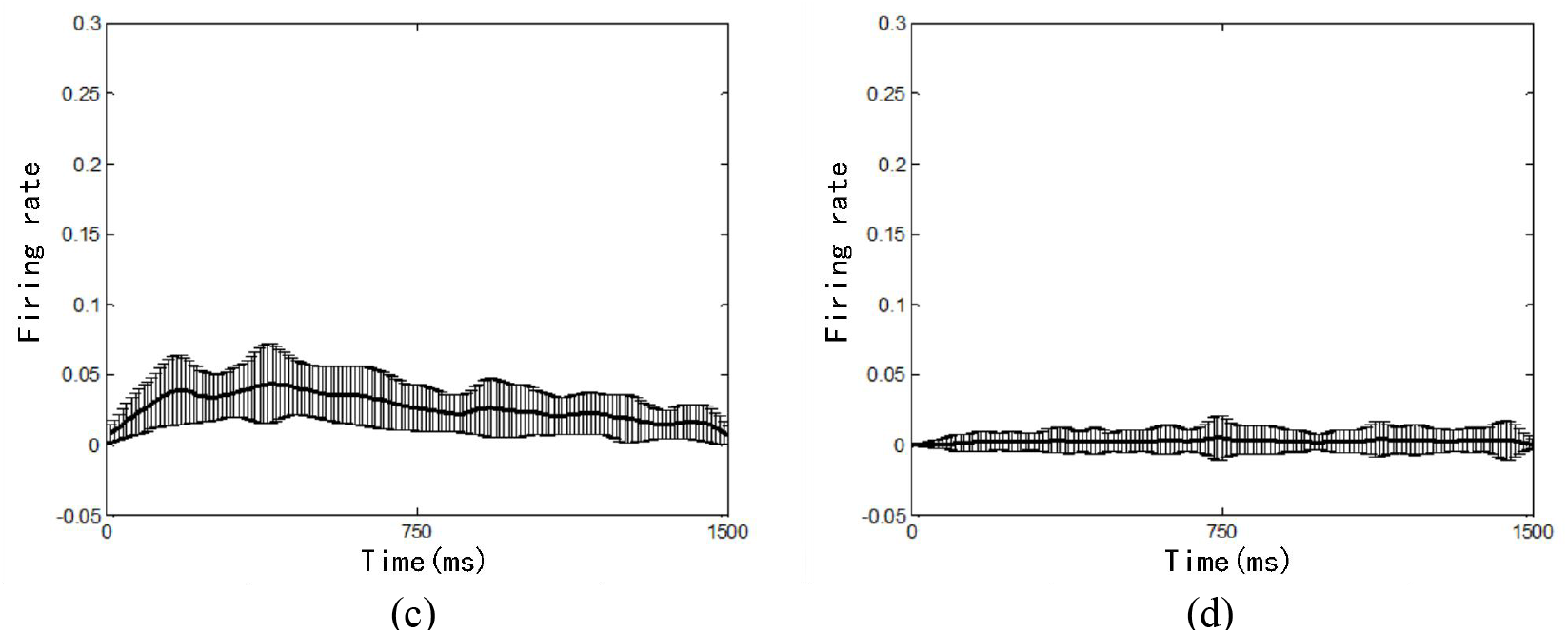
The statistical response result under continuous sensitivity stimuli.The thick solid line represents the mean value of firing rate, and the thin solid line represents its corresponding variance. The average curves were smoothed by Gauss filter. (a) and (c) are the electrophysiological experiment statistical result to random sequential motion (1/4) and random motion using 14 collected continuous motion-sensitive neurons. (b) and (d) are model statistical result to random sequential motion (1/4) and random motion using 300 simulated continuous motion-sensitive neurons which were randomly generated and had somas located in one same center of the receptive field.

Since these continuous motion-sensitive neurons with different sizes of receptive fields showed consistent statistical patterns, some simulate continuous motion-sensitive neurons which had fixed-size of receptive fields (18°×18°) were used to observe the model response patterns to continuous sensitivity stimuli. The tuning property and the statistical response result of model to continuous sensitivity stimuli was shown in Fig. 13 (black line) and Fig. 12b and 12d. The variation trend of variance both in Fig. 12a and 12b can be clearly seen that the whole path was divided into four continuous parts for random sequential motion (1/4). The statistical firing rate of model was lower than that of the physiological experiments for random motion. The tuning property of the model is basically consistent with the overall trend of the tuning property of electrophysiology, which again proves the validity of our model and hypotheses. Random sequential motion can be regarded as the transition state of sequential motion and random motion. It is clear that continuous motion-sensitive neurons prefer motion with high continuity. The longer local sequential motion length of the random sequential motion (the larger the rate of the path length), the more likely the activated dendrites are to accumulate directed energy *E*, and the more likely the activated dendrites to induce somatic response. The shorter local sequential motion length of the random sequential motion (the smaller of the rate of path length), the more discrete the track is (the extreme case is random motion), and the more likely it is to lose directed energy *E*, leading to induce weaker somatic response.

**Fig.13.**
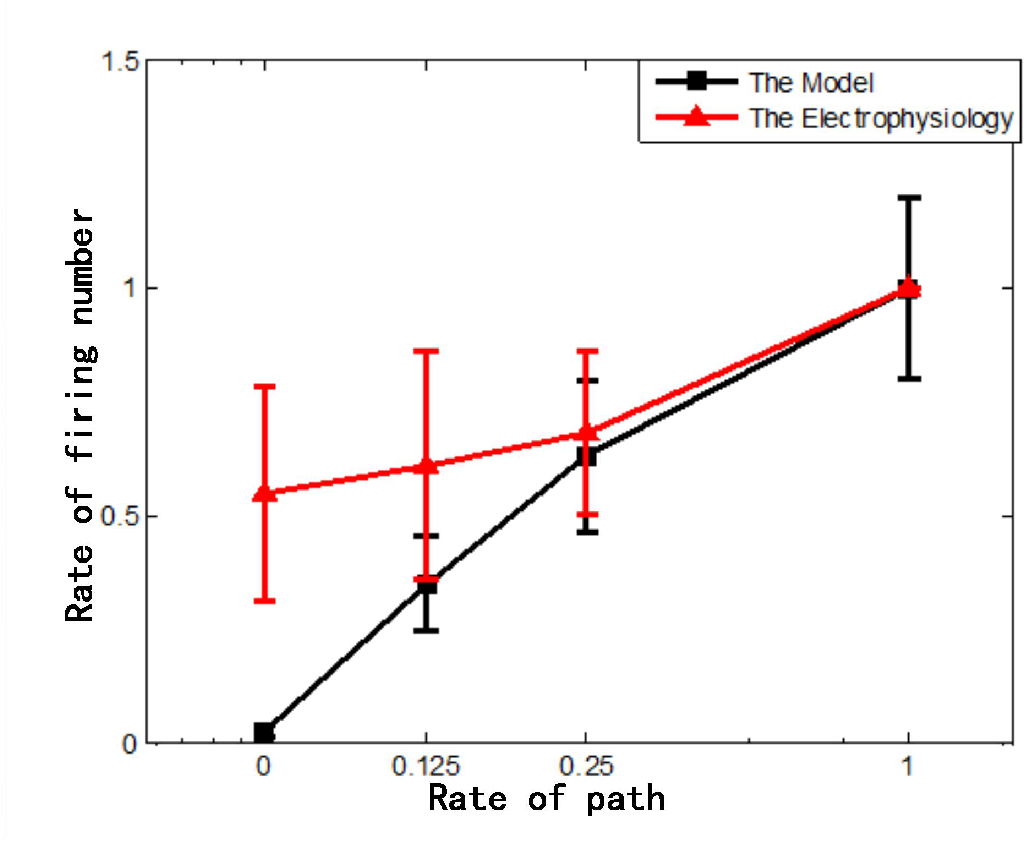
The tuning curves of continuous motion-sensitive neurons to continuous sensitivity stimuli.The red triangle line represents the result produced by the electrophysiology using 14 collected continuous motion-sensitive neurons. The black square line represents the result produced by model under using 300 simulated continuous motion-sensitive neurons. The firing number of each neuron for each kind of motion was calculated, and then divided by the firing number of sequential motion to normalize. The firing rate of all neurons was then added up and normalized again. The horizontal axis is the logarithmic axis and represents the rate of path lengh (random motion, random sequential motion (1/8), random sequential motion (1/4), sequential motion), and the vertical axis represents the mean value and variance of rate of firing number.

## 4 Discussion

In this paper, we implanted OT neurons at depths of 800-1500μm to study their encoding model of small target in continuous motion. These continuous motion-sensitive neurons prefer small continuous moving targets and their single dendrite activation can induce somatic response. Based on the above phenomena, the innovative directed energy accumulation hypothesis in dendrite field and sequential probability activation hypothesis of soma were proposed. Then, an information encoding model of these kind of neurons is constructed based on the proposed hypotheses. Physiological experiments and special stimuli furthermore helped us to validate qualitatively the validity of the model.

In order to model the preference of continuous motion-sensitive neurons for continuous motion, the directed energy accumulation hypothesis is proposed in this paper. This hypothesis assumes that the activated dendrite will pre-activate an area along the direction of its predicted motion track, which will enhance the ability of the subsequent activated dendrite in this area to induce the somatic response. Although the model based on this hypothesis has obtained acceptable model fitting results, there is no direct experiment to provide some plausible evidences for this hypothesis. Here, we will discuss some of the grounds for giving this hypothesis.

The highly structured sensory signals in both space and time for visual information allow expectations about future stimulation to facilitate sensory processing and decision-making (Summerfield & De Lange,2014). Perceptual prediction is usually studied in static settings where the stimulus is expected because of the higher base incidence (Squires.et al.,1976) or because of the statistical association between the stimuli (Meyer.et al.,2014). These forms of prediction can be neurologically realized by preactivating sensory representations of the expected event (Esterman & Yantis,2009; Sakai & Miyashita,1991; Schlack & Albright,2007). However, given the dynamic nature of prediction in the real world: for example, we predict the track of a ball moving towards us, implementing this kind of dynamic prediction is more complex, as it requires an anticipatory wave of visual response that is both spatially and temporally precise. Recently, such waves of preplay activity have been observed in the visual cortical system of mice (Xu & Jiang,2012) and monkeys (Eagleman & Dragoi,2012), and even humans (Ekman.et al.,2017).

In (Ekman.et al.,2017), they found that flashing only the starting point of a moving dot sequence triggered an activity wave in V1 that recreates the full stimulus sequence. This anticipatory activity wave was temporally compressed compared to the actual stimulus sequence and was present even when attention was diverted from the stimulus sequence. This preplay activity may reflect an automatic prediction mechanism for visual sequences. This means that when neurons do not receive the physical stimulation of the moving target, they can achieve the dynamic prediction of the moving target through sequential activation at different moments in the future, so as to facilitate subsequent related actions and behaviors. But how this “rehearsal” facilitates subsequent action remains unclear. The directed energy accumulation hypothesis proposed in this paper can be regarded as a realization of the facilitation of this “rehearsal” for future target detection.

The sequential probabilistic activation hypothesis is supported by solid experimental phenomena in both avian and mice.

In (Luksch et al., 2004), it was found that as long as the time interval of continuous bbe activation was greater than a certain threshold, a single activated bbe would induce the somatic response with a high probability. In detail, they locally stimulated a small group of RGC axons with short current pulses that were delivered with a stimulus electrode in layers 2-4 and recorded the response in the soma of continuous motion-sensitive neurons to investigate the signal transfer from the RGC axon to the soma of continuous motion-sensitive neurons. In all continuous motion-sensitive neurons testes, single-pulse stimulation resulted in an all-or-none sharp-onset somatic response consisting of either one to three action potentials riding on a broader depolarization or no response. Then, they carried out one-site regular pulse train synaptic stimulation experiments. The sharp-onset response to each stimulus pulse was probabilistic. For all stimulation intervals tested, the response probability reached a steady state after the second stimulus pulse. In other words, for stimulation at the same site (alone) on RGC, with the increase of time interval between stimuli, the probability of continuous induction of somatic response by a single dendrite increases continuously. To investigate the cell’s response to spatiotemporal synaptic inputs and to test for potential distance dependence of the interaction, they placed two stimulus electrodes in layers 2-4 at distances of 250-1,500 µm apart, thus stimulating two separate groups of RGC axons, and recorded from a continuous motion-sensitive neuron that received inputs from both groups of axons and stimulated the two sites in temporal sequence with varying stimulation intervals. The measured response probabilities to the second stimulus pulse for one stimulation interval showed no statistically significant distance dependence for the two-site synaptic stimulation. In particular, when the time interval between successive stimuli at two different locations on the RGC exceeds 100ms, the probability of stimulating somatic response on a single dendrite is almost 1.

As for the mice SC, Gale showed that input restricted to a small portion of the broad dendritic arbor of WF cells is sufficient to trigger dendritic spikes that reliably propagate to the soma/axon (Gale and Murphy, 2016). In vivo whole-cell recordings reveal that nearly every action potential evoked by visual stimuli has characteristics of spikes initiated in dendrites. Moreover, to verify whether previous bouts of synaptic input influence the ability of WF cell dendrites to generate spikes, a broad swath of ChR2-expressing retinal axons was simultaneously activated by light pulses delivered through the microscope objective. The results shown that dendritic spikes were reliably evoked by sequential light pulses separated by as little as 20ms in all WF cells. In other words, the single dendrite of mice WF neuron has the ability to induce the somatic response, but the conclusion on the probability of the single dendrite inducing the somatic response is not the same as that of avian. In continuous motion-sensitive neurons of the avian optic tectum, one-site regular pulse train was stimulated at a small group of RGC axons and the dendritic endings respectively, which shows that the somatic spikes can be stably induced at both locations at a certain pulse interval and signal transfer within continuous motion-sensitive neurons is tonic at time scales that are two orders of magnitude shorter than signal transfer at the retinotectal synapse. So, strong and prolonged (lasting seconds) synaptic inhibition was proposed to underlie phasic signal transduction at inputs to an individual dendritic branch and therefore selective response to stimuli that move across several branches (Luksch et al.,2004).

In general, a single dendritic possess the ability to induce somatic response both for the avian continuous motion-sensitive neurons and WF cells in mice, but the probability of inducing somatic response by dendritic did not show a clear guideline. Considering the motion preference of this kind of neurons show, the hypothesis proposed in this article that the probability depends on the energy value of dendrites which is a reflection of the continuity of dendritic sequential activation has the biological plausibility.

Since Hubel and Wessle discovered neuronal orientation selectivity from the primary visual cortex of cats in 1962 (Hubel D H and Wiesel T N, 1962), biological and neuroscience research on the analysis of the information processing mechanism of the mammalian retina-primary visual cortex-inferior temporal lobe of the hypothalamic visual pathway, and the interaction of information science research that uses various mathematical methods to describe and model related neural mechanisms, have greatly promoted the progress of their respective fields, and at the same time these progress have brought huge enlightenment to the task of target recognition in machine vision. Its representative is deep learning, convolutional neural networks and other algorithms with far-reaching influence in the recent years. Avian have excellent visual perception at high-altitude and high-speed. The tectofugal visual pathway of avian more which developed than mammals is the neural basis for realizing this visual perception ability. Therefore, analyzing the information processing mechanism of the typical continuous motion-sensitive neurons to weak and small targets in the OT of avian and constructing an encoding model will provide inspiration for the formation of a new brain-like target detection system that is different from the existing deep learning.

## Notes

### Competing Interest Statement

The authors have declared no competing interest.

## References

Drager U C, Hubel D H. 1975. Responses to visual stimulation and relationship between visual, auditory, and somatosensory inputs in mouse superior colliculus[J]. Journal of Neurophysiology, 38(3):690–713.

DeAngelis G, Ohzawa I, & Freeman R.D. 1995. Receptive-field dynamics in the central visual pathways. Trends in Neurosciences, 18, 451–458.

Diggle PJ. 2003. Statistical analysis of spatial point patterns. London. Arnold.

Endo, T, Yanagawa, Y, Obata, K, & Isa, T. 2003. Characteristics of gabaergic neurons in the superficial superior colliculus in mice. Neuroscience Letters, 346(1-2), 81–84.

Endo T. Tarusawa, E, Notomi, T, Kaneda, K, Hirabayashi, M, & Shigemoto, R, et al. 2008. Dendritic ih ensures high-fidelity dendritic spike responses of motion-sensitive neurons in rat superior colliculus. Journal of Neurophysiology, 99(5), 2066–2076.

Esterman, M, & Yantis, S. 2009. Perceptual expectation evokes category-selective cortical activity. Cerebral Cortex, 20(5), 1245–1253.

Eagleman, S. L, & Dragoi, V. 2012. Image sequence reactivation in awake v4 networks. Proceedings of the National Academy of Sciences, 109(47), 19450–19455.

Frost, B. J, & Difranco, D. E. 1976. Motion characteristics of single units in the pigeon optic tectum - sciencedirect. Vision Research, 16(11), 1229–1234.

Frost, B. J. 1993. Subcortical analysis of visual motion: relative motion, figure-ground discrimination and self-induced optic flow. Rev Oculomot Res, 5(5), 159–175.

Hubel, D. H, & Wiesel, T. N. 1962. Receptive fields, binocular interaction and functional architecture in the cat’s visual cortex. Journal of Physiology, 160(1), 106–154.

Harald, Luksch, Kevin, Cox, Harvey, & J., et al. 1998. Bottlebrush dendritic endings and large dendritic fields: motion-detecting neurons in the tectofugal pathway. The Journal of Comparative Neurology.

Jassik-Gerschenfeld D, Guichard J, Tessier Y. 1975. Locialization of directionally selective and movement sensitive cells in the optic tectum of the pigeon.[J]. Vision Research, 15(8-9):1037, IN11-1038,IN11.

Jan, Y, & Jan, L. 2003. The Control of Dendrite Development. Neuron, 40, 229–242.

Karten, H, Cox, K, & Mpodozis, J. 1997. Two distinct populations of tectal neurons have unique connections within the retinotectorotundal pathway of the pigeon (Columba livia). Journal of Comparative Neurology, 387(3), 449–465.

Khanbabaie, R, Mahani, A. S, & Wessel, R. 2007. Contextual interaction of gabaergic circuitry with dynamic synapses. Journal of Neurophysiology, 97(4), 2802–2811.

Luksch H, Khanbabaie R, Wessel R. 2004. Synaptic dynamics mediate sensitivity to motion independent of stimulus details.[J]. Nature Neuroscience, 7(4):380.

Luksch, H, & Golz, S. 2003. Anatomy and physiology of horizontal cells in layer 5b of the chicken optic tectum. Journal of Chemical Neuroanatomy, 25(3), 185–194.

Luksch, H, Cox, K, & Karten, H. 1998. Bottlebrush dendritic endings and large dendritic fields: Motion-detecting neurons in the tectofugal pathway. Journal of Comparative Neurology, 396.

Mahani, A. S, Khanbabaie, R, Luksch, H, & Wessel, R. 2006. Sparse spatial sampling for the computation of motion in multiple stages. Biological Cybernetics, 94(4), 276–287.

Meyer, T, Walker, C, Cho, R. Y, & Olson, C. R. 2014. Image familiarization sharpens response dynamics of neurons in inferotemporal cortex. Nature Neuroscience, 17(10), 1388–1394.

O’Leary D D M, Mclaughlin T. 2005. Mechanisms of retinotopic map development: Ephs, ephrins, and spontaneous correlated retinal activity[J]. Progress in Brain Research, 147:43–65.

Squires, K, Wickens, C, Squires, N, & Donchin, E. 1976. The effect of stimulus sequence on the waveform of the cortical event-related potential. Science, 193(4258), 1142–1146.

Sakai, K, & Miyashita, Y. 1991. Neural organization for the long-term memory of paired associates. Nature, 354(6349), 152–155.

Stuart G, Spruston N, Haeusser M. 1999. Dendrites. Oxford. Oxford University Press.

Schlack, A, & Albright, T. D. 2007. Remembering visual motion: neural correlates of associative plasticity and motion recall in cortical area mt. Neuron, 53(6), 881–890.

Summerfield, C, & De Lange, F. P. 2014. Expectation in perceptual decision making: neural and computational mechanisms. Nature Reviews Neuroscience, 15(12), 816–816.

S. D, Gale, G J, & Murphy. 2014. Distinct representation and distribution of visual information by specific cell types in mouse superficial superior colliculus. Journal of Neuroscience, 34(40).

S, D, Gale, G, J., & Murphy. 2016. Active dendritic properties and local inhibitory input enable selectivity for object motion in mouse superior colliculus neurons. Journal of Neuroscience.

Verhaal, J, & Luksch, H. 2016. Neuronal responses to motion and apparent motion in the optic tectum of chickens. Brain Research, 1635, 190–200.

Wang, S, Wang, M, Wang, Z., & Shi, L. 2019. First spike latency of on/off neurons in the optic tectum of pigeons. Integrative Zoology, 14(5).

Xu, S, Jiang, W, Poo, M, & Dan, Y. 2012. Activity Recall in Visual Cortical Ensemble. Nature neuroscience, 15, 449–452.

